# Chronological aging impacts abundance, function and microRNA content of extracellular vesicles produced by human epidermal keratinocytes

**DOI:** 10.1101/2022.10.31.514487

**Authors:** Taku Nedachi, Christelle Bonod, Julie Rorteau, Wafae Chinoune, Yuri Ishiuchi, Sandrine Hughes, Benjamin Gillet, Dominique Sigaudo-Roussel, Jérôme Lamartine

**Affiliations:** Skin Functional Integrity group. Laboratory for Tissue Biology and Therapeutics Engineering (LBTI) CNRS UMR5305 – University of Lyon – France; Department of Life Science – Toyo University – Japan; IGFL CNRS UMR5242 – ENS de Lyon – University of Lyon – France

## Abstract

The disturbance of intercellular communication is one of the hallmarks of aging. The goal of this study is to clarify the impact of chronological aging on extracellular vesicles (EVs), a key mode of communication in mammalian tissues. We focused on epidermal keratinocytes, the main cells of the outer protective layer of the skin which is strongly impaired in the skin of elderly. EVs were purified from conditioned medium of primary keratinocytes isolated from infant or aged adult skin. A significant increase of the relative number of EVs released from aged keratinocytes was observed whereas their size distribution was not modified. By small RNA sequencing, we described a specific microRNA (miRNA) signature of aged EVs with an increase abundance of miR-30a, a key regulator of barrier function in human epidermis. EVs from aged keratinocytes were found to be able to reduce the proliferation of young keratinocytes, to impact their organogenesis properties in a reconstructed epidermis model and to slow down the early steps of skin wound healing in mice, three features observed in aged epidermis. This work reveals that intercellular communication mediated by EVs is modulated during aging process in keratinocytes and might be involved in the functional defects observed in aged skin.

## Introduction

The most obvious sign of the aging process is skin aging. Because of its accessibility and because subjected to environmental variations and stresses, the skin is an ideal organ for investigating the different components of aging and its underlying causalities in humans. Skin aging is a complex biological process involving a combination of intrinsic (endogenous) and extrinsic factors. During this natural aging process, the skin undergoes a physiological deterioration characterized by skin atrophy, increased physical and immunological vulnerability, with a reduced capacity of tissue repair in case of wounding. Extrinsic aging caused by external factors such as sun exposure or smoking may prematurely age the skin. The epidermis, the outer layer of the skin, is especially confronted with the various components of aging causes. Aged human epidermis displays several morphological changes, such as the decreasing in epidermal thickness and flattening of the dermal-epidermal junction. Some of these changes can be explained by a decrease in proliferation, an impairment of differentiation and the induction of apoptosis in keratinocytes, that represent the main epidermal cellular population [1, 2]. Several cell-autonomous mechanisms have been proposed to be involved in this aging process such as dysfunction of mitochondria, shortening of telomeres, loss of proteostasis or genomic instability, but changes at the level of intercellular communication represents also an important hallmark of aging [3]. Moreover, cells with intracellular aged-related defects might be able to propagate a deleterious message to the surrounding cells and tissues. This has been observed for senescent cells, which critically modulate their extracellular environment notably through the secretion of pro-inflammatory cytokines and metalloproteases, and then contribute to the propagation of the aging phenotype [4].

Intercellular communications can be achieved via at least three distinct mechanisms: secretion of small molecules, direct cell-cell contact and release of extracellular vesicles (EVs). The two first mechanisms have been extensively described, notably in senescent skin fibroblasts with the secretion of pro-inflammatory cytokines and the transmission of a senescence signal related to neighboring cells via gap junction [5, 6]. Nevertheless, among these mechanisms, EVs-mediated intercellular communication has recently accumulated much attention, since numerous studies demonstrated tight associations between the release of EVs and many physiological and pathological processes, especially in skin [7]. It has been indeed observed that EVs are involved in the cutaneous immune response, skin pigmentation and wound healing, but also in skin diseases such as melanoma and psoriasis [7]. EVs are divided into 3 different groups with various sizes: apoptotic bodies (with a diameter comprised between 1 and 5 μm), microvesicles formed by direct budding of the plasma membrane (from 0.2 to 2 μm) and exosomes (from 50 to 150 nm) that are originated from multivesicular bodies containing intraluminal vesicles that are trafficked to the cell surface in a Rab-27 dependent manner [8]. EVs carry a complex cargo containing not only lipids, but also many types of proteins and nucleic acids. Strikingly, the recent findings that various RNAs notably mRNAs, miRNAs, lncRNas and snoRNAs are present in EVs strongly suggest that these RNAs play a central role in transferring signal from one cell to another. We observed that miRNA expressional profiles in human keratinocytes are dramatically changed during the chronological aging process [9]. It can be thus presumed that these changes may directly influence the miRNA profiles in EVs released from these keratinocytes and that a specific signal might be transmitted through EVs by aged cells. However, to the best of our knowledge, this hypothesis remains unexplored, at least in skin.

The present study was therefore conducted to elucidate whether the characteristics of EVs released from cultured human keratinocytes can be modulated during aging process. We carried out an exhaustive inventory of the miRNAs present in EVs secreted by keratinocytes prepared from young or old skin. We also compared the functional impact of the EVs released from keratinocytes prepared from young or aged skin samples. Altogether, our results show that chronological aging modulates abundance and miRNAs content of keratinocytes EVs. Concurrently, we observed that the EVs extracted from aged cells exerted an anti-proliferative and anti-migratory effect on young keratinocytes *in vitro* and *in vivo*. This suggest that aged cells might modulate the function of their surrounding cells through EVs secretion.

## Materials and methods

### Ethical consideration and cell culture

Normal human skin tissue explants were obtained from the surgical discard of anonymous healthy patients with informed consent of adult donors or children’s parents in accordance with ethical guidelines (French Bioethics law of 2004) and declared to the French research ministry (Declaration no. DC-2008-162 delivered to the Cell and Tissue Bank of Hospices Civils de Lyon). Young (<5 years), and aged (>60 years) human primary keratinocytes (HPK) were isolated with trypsin (Gibco, Life Technologies, Carlsbad, CA, USA) and dispase (Dispase II; Roche Diagnostics, Mannheim, Germany), respectively from child foreskin or abdominal biopsies obtained from plastic surgery. Keratinocytes isolation, culture in keratinocyte growth medium 2 (KGM2; PromoCell, Heidelberg, Germany) were performed as previously described [10]. Cells were seeded on appropriate flasks (Falcon) in KGM2 medium (PromoCell) supplemented with 100 μg/ml Primocin™ (InvivoGen). Subconfluent cultures were passed using 0.05% trypsin/EDTA (GIBCO). For young or aged conditions, 3 independent culture of primary keratinocytes were established from 3 independent biospies.

### EVs isolation

Extracellular vesicles were prepared as described previously [11]. Briefly, subconfluent keratinocytes were continuously cultured for 48h and conditioned media were collected. Media were centrifuged at 2000g for 15 min and 4000g for 15 min to remove cell debris. The supernatant was then centrifuged at 10000g for 30 min, and the obtained supernatant were subjected to final ultracentrifugation at 100000g for 60 min. The pellets were washed with PBS and resuspended in 1 ml of PBS. Finally, EVs suspension was subjected to ultrafiltration (3KMO) to obtain concentrated EVs.

### Protein analysis

Keratinocytes or EVs were lysed with radio-immunoprecipitation assay (RIPA) buffer (10 mM Tris pH8.8, 150 mM NaCl, 1 mM EDTA, 1% NP40, 1% sodium deoxycholate, 0.1% SDS). Protein concentration in each sample was measured by BCA protein assay kit (Pierce). Protein samples were then subjected to SDS-PAGE (10%) and visualized by silver staining (Pierce) or transferred to PVDF membranes (Millipore). The membrane was blocked in TBS/Tween 0.1% (TBS-T) with 5% non-fat skim milk, incubated with the indicated antibodies anti-CD63 (Santa Cruz), anti-TSG101 (System Biosciences), anti-GM130 (AbCam), anti-p16 and anti-IDH1 (1/1000, Cell Signaling Technology) and anti-actin (1/500, Millipore) for overnight. Specific total proteins were visualized after subsequent incubation with a 1/10000 dilution of antimouse, anti-rabbit, or anti-goat IgG conjugated to horseradish peroxidase and a SuperSignal Femto Chemiluminescence detection procedure (Pierce Biotechnology Inc.). At least three independent experiments were performed for each condition.

### Dynamic light scattering analysis for EVs

Hydrodynamic diameter of EVs and size distribution were estimated by dynamic light scattering (DLS) using a Zetasizer NanoZSP (Malvern Instruments, U.K.). EVs in PBS buffer were analyzed in a 10 mm path length cuvette at a temperature of 25°C and a scattering angle of 173°. Each value was the average of four series of 10 measurements.

### Proliferation and migration assays

Proliferation was monitored using the CyQUANT Cell Proliferation Assay kit (Thermofischer Scientific) following the manufacturer instructions. Keratinocytes were seeded in a 96 wells plate at 3000 cells/well. The DNA quantification test was performed after 2h, 48h and 72h of incubation with 2 μg of EVs extracted from conditioned medium of young or old keratinocytes. For studying the migration of keratinocytes, a scratch wounding assay was performed. Keratinocytes were seeded at 70000 cells/well in a 12 wells plate. After cell reach 100% confluence, a wound was made in each well using a tip200. Wells were rinsed 2 times with PBS and new KGM2 medium was added supplemented with 1μg of old exosomes. Wound filling was monitored by light microscopy 24h and 40h after wounding. The surface of remaining wound was measured using ImageJ.

### Culture of reconstructed human epidermis (RHE)

RHE preparation was performed exactly as previously described [9] using keratinocytes from infant foreskin or from aged abdominal skin (> 60 years old) and fibroblasts from adult skin. Briefly, keratinocytes were seeded at 300000 cells per insert. After an initial incubation of 72h in DMEM-F12 without vitamin C, the culture was kept emerged at the air liquid interface for 14 days. The medium (DMEM-F12 + CaCl2 supplemented with vitamin C) was changed daily and supplemented with EVs (0.25 μg per ml) every 2 days. The RHE were finally fixed with Zinc Formal-FIxx (ThermoFischer Scientific) at 4°C overnight and embedded in paraffin. The sections were stained with hematoxylin, eosin and safranin or used for immmuno-histochemistry.

### Wound healing assay in mice

All experiments were performed following the principles of French legislation and were approved by the Ethics Committee of Animal Experiments of Lyon. 12-week-old male C57BL/6J mice (Charles River laboratories, l’Arbresle, France) were allowed to adapt to their environment for 7 days before the experiments. Mice were maintained at 21°C on a 12h light/dark cycle with food and water *ad libitum*. All efforts were made to minimize suffering.

Mice were housed in individual cages after wounding. Mice were anesthetized with isoflurane and after shaving the dorsal hairs and disinfecting the skin, a 8-mm diameter full-thickness wound was created on the middle back. Mice were randomly divided into two groups. Exosomes-group (8 animals): 4 intradermal injections were carried out at 4 diametrically opposite points (12.5 μg of EVs in 50 μl in each injection) around the punch. Control group (4 animals): injection of NaCl 0.1% instead of exosomes. Pictures were taken every day until closure. The area of the wound was quantified using ImageJ and expressed as a percentage of the initial wound area. Mice were killed at the end of the healing and wound skin tissues were collected for histology and immunofluorescence staining.

### Quantitative PCR analysis

Total RNA from cells or EVs were purified by using MirVana miRNA isolation kits (Thermofisher). For evaluating gene expression in cells, the first-strand cDNA was synthesized by using PrimeScriptTM RT reagent kit (Takara). Quantitative PCR analysis was performed with SYBR Premix Ex Taq II on AriaMx Real-time PCR system (Agilent). Specific primer sets for age-related genes and housekeeping gene were used: p16 F: 5’- GAAGGTCCCTCAGACATCCC-3’; p16 R: 5’-CCCTGTAGGACCTTCGGTGA-3’, IDH1 F: 5’-TTGGCTGCTTGCATTAAAGGTT-3’; IDH1 R: 5’-GTTTGGCCTGAGCTAGTTTGA-3’, TBP F: 5’-TCAAACCCAGAATTGTTCTCCTTAT-3’; TBP R: 5’-CCTGAATCCCTTTAGAATAGGGTAGA-3’.

### Small RNA libraries preparation

12 total RNA samples extracted from keratinocyte small EVs (Exosomal RNA from young keratinocyte early passage referenced 025 P2, Y205 P2, Y215 P2 and Y220 P2, exosomal RNA from old keratinocyte early passage referenced A12015 P2, A11162 P2, A059 P and A175 P2, exosomal RNA from young keratinocyte late passage referenced Y025 P6, Y205 P6, Y215 P6 and A Y220 P6). First, they were concentrated from 40 μl to 5 μl during 2h at low temperature using speedvac (Thermofisher). Small RNA libraries were then built with Illumina TruSeq Small RNA Sample kit following manufacturer’s recommendations. 15 PCR cycles were conducted for libraries amplification. Each library was size-selected by running them in a 6% TBE Gel (Thermofisher) for 60 min at 145 V. After staining the gel with ethidium bromide, the gel band between 140bp to 160bp were cut using a razor blade. They correspond to the adapter-ligated constructs derived from the 22 nt and 30 nt small RNA fragments. Gel pieces were crushed by centrifugation at 20000g for 2 min using Gel Breaker tubes (IST Engineering BioTech). Gel debris were incubated in 200 μl of 0.5 M Sodium Acetate, 10 mM Magnesium Acetate, 1 mM EDTA pH8, 0.1% SDS for 2h at 50°C with shaking at 500rpm using Thermomixer (Eppendorf). Then, the eluate and gel debris were transferred to the top of a 5 μm filter (IST Engineering BioTech) and centrifuged 10 sec at 600g. Finally, the llumina size-selected libraries were converted by PCR to be sequenced on Ion Torrent Proton sequencer (Thermofisher). PCR mix conditions used were: 25 μl at 400 pM of Illumina size-selected library, 100 μl Platinum PCR SuperMix High Fidelity (Thermofisher), 2.5 μl primer fusion A 10μM 5’-CCATCTCATCCCTGCGTGTCTCCGACTCAGXXXXXXXXXXGATTCCCTACACGAC GCTCTT-3’ (X corresponds to the Ion Torrent 10 long nucleotides barcode, numbers 23, 24, 25 and 26 were used in this study, respectively TGCCACGAAC, AACCTCATTC, CCTGAGATAC and TTACAACCTC) and 2.5 μl fusion primer P1 10 μM 5’-CCTCTCTATGGGCAGTCGGTGATGAGTTCTGACGTGTGCTC-3’. PCR cycling conditions were: 95°C for 5 min, followed by 8 cycles of 95°C for 15 sec, 58°C for 15 sec, and 70°C for 1 min. Amplified converted libraries were purified using SPRI paramagnetic bead (Agencourt AMPure XP, Beckman Coulter) with a ratio of beads/DNA in solution of 1.2X and eluted in 25 μl of Low TE. Quantitation and quality assessment of each of the 12 libraries were performed on 2200 Tapestation analyzer using the High Sensitivity D1000 ScreenTape kit (Agilent).

### Small RNA sequencing

Libraries were sequenced using Proton sequencer at the Sequencing Platform of the IGFL. To increase read numbers, four sequencing runs were done using 3 libraries mixed in an equimolar manner for a run. Template preparation (emulsion PCR, Ion Sphere particles enrichment) and PI chip loading were conducted using the Ion Chef System (Thermofisher) with an input of 25 μl of a 75 pM equimolar libraries mix. Sequencing runs were conducted on Proton sequencer using the PI™ Hi-Q™ Sequencing 200 Kit (Thermofisher) following the recommendations of the manufacturer. They allowed to obtain from 90M to 127M of reads per run and 24M to 40M of reads per library.

### Small RNAseq data analysis

Quality of raw reads was assessed using the FastQC quality control tool. The various RNA species present in the samples were quantified starting from the raw read data (fastq files) using miRge [12]. miRNA quantification was also performed using Cutadapt to remove the adapters, followed by functions qAlign and qCount from package QuasR. For the quality control representations, the effective library sizes were first computed using function estimateSizeFactors from DESeq2 package. The raw count values were then transformed using a variance stabilizing transformation (function variance Stabilizing Transformation from package DESeq2). miRNA that did not have more than one count per million counts in at least 3 samples were filtered out. Hierarchical clustering of samples was performed using Ward’s agglomerative method, passing the euclidean distances between samples to function hclust from package stats. Cluster stability was estimated by multiscale bootstrap resampling using function pvclust from pvclust R package. Hierarchical clustering of miRNA was performed using the complete agglomerative method, passing the Euclidean distances between centered (for expression data) or uncentered (for fold change data) and scaled signals to function hclust from R Stats package. Principal Component Analyses (PCA) of sample expression levels was performed with miRNA signals centered but not scaled (using function prcomp from package Stats). When displayed, miRNA coordinates correspond to these miRNA correlations with the presented components. Only miRNA that contribute most to the components are displayed. Differential expression was assessed using the limma package. The empirical Bayes method (function eBayes from the limma package) was used to compute moderated *p*-values. *p*-values were then corrected for multiple comparisons using the Benjamini and Hochberg’s false discovery rate (FDR) controlling procedure.

### Statistical analysis

All statistical analyses for zetasizers, QPCR, proliferation and migration data were performed with Prism software (GraphPad Software Inc., La Jolla, CA, USA). Results are expressed as mean value ± SEM, and the data were analyzed by student t-test or the one-way analysis of variance (ANOVA) followed by Tukey’s posthoc test. Differences were considered to be significant at *p < 0.05; **p < 0.01; and ***p < 0.001

## Results

### EVs purification from human keratinocyte culture

In order to purify EVs from keratinocytes conditioned media, we used a classical ultracentrifugation method, which consists of several centrifugation steps aiming to remove cell debris and precipitate EVs as described in Materials and Methods. A protein lysate of purified EVs was obtained and analyzed for the expression of cellular fraction markers: the expression of exosomal markers CD63 and TSG101 was clearly confirmed by Western-blot analysis, whereas no detectable expression of the golgi marker protein, GM130 was observed (Fig. 1A). On the other hand, all of these protein expressions were clearly observed in whole cell lysates including GM130 (Fig. 1A). Electron transmission microscopic analysis revealed the cupshaped vesicles that are typically observed in negative staining of EVs [13] and that the diameter was approximately 50 nm, which suggested the purified EV fraction contained a population of small vesicles (Fig. 1B). Moreover, dynamic light scattering (DLS) analysis also suggested the population of vesicles was distributed around 50-200 nm in diameter, and z-average of vesicles was 154.367 ± 3.197 d. nm (Fig. 1C). The polydispersity index (PDI) of samples was relatively high (0.324 ± 0.034), suggesting that purified EVs were either heterogeneous or aggregated at some extent. In addition to this intensity-based size distribution, we also analyzed number- and volume-based size distribution (Fig. 1C). Overall, these results indicate that the classical ultracentrifugation method allowed to purify EVs from keratinocyte culture supernatant and this purified fraction contained mostly exosomes, based on Western-blot analysis, even though we could not exclude the possibility of the presence of small microvesicles. Since currently available purification methods do not allow us to discriminate perfectly between exosomes and MVs [14], we called this fraction as “small EV fraction” and used them for the following experiments.

**Figure 1:**
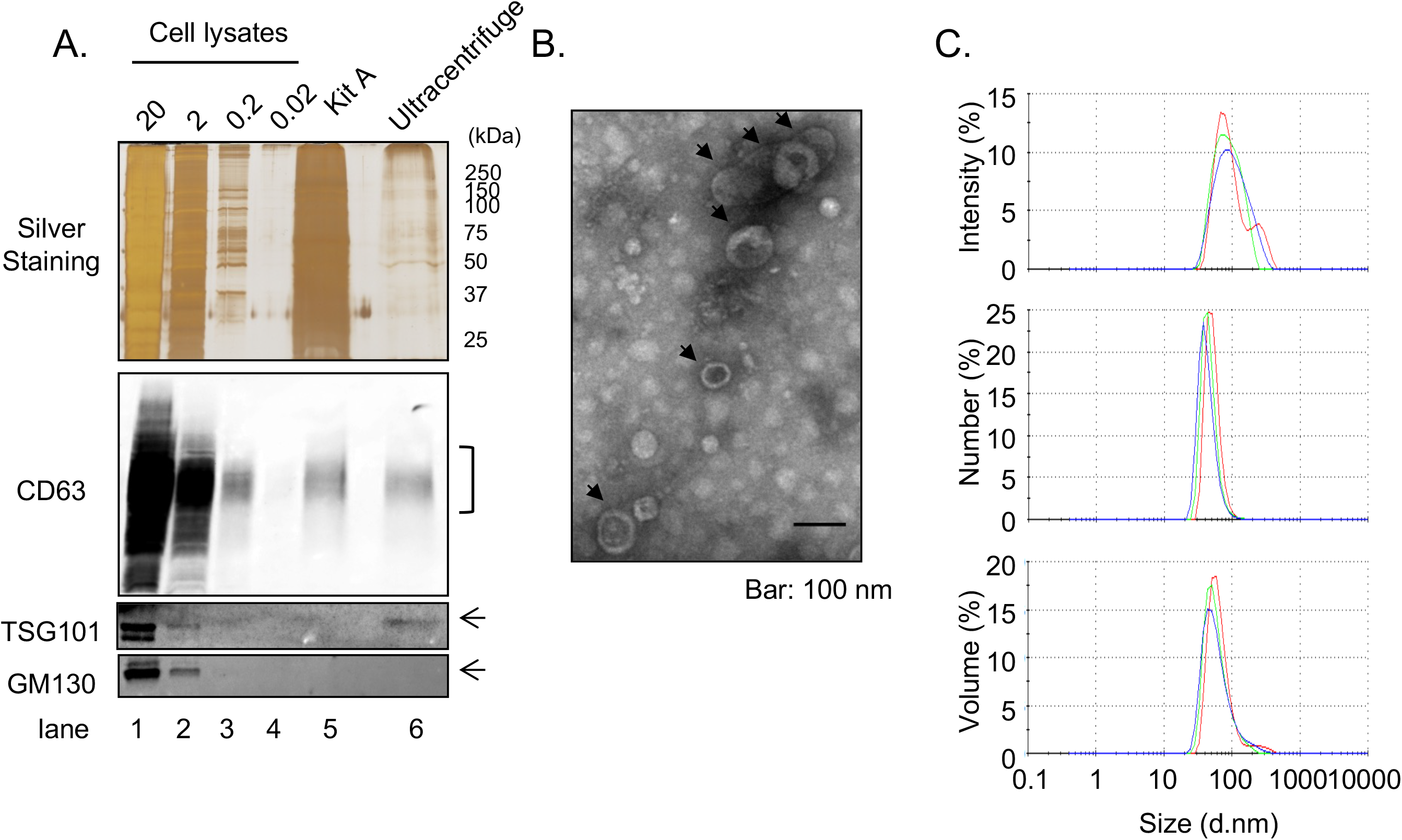
Isolation of EVS from cultured keratinocytes. Conditioned media from human primary keratinocytes were collected. EVs were isolated from these media by an ultracentrifugation protocol. (A) The expression of specific markers for EVs (CD63 and TSG101) were observed in the ultracentrifugation purified EVs fraction (Ultracentrifuge lane), whereas the expression of GM130 were not observed. 20 μg, 2 μg, 0.2 and 0.02 μg of total cell lysates were also analyzed as well as an EV fraction purified by the Qiagen exoEasy kit (Kit A lane) (B) Analysis of ultracentrifuge purified EVs fraction by transmission electronic microscopy. (C) Hydrodynamic size distribution profiles of ultracentrifuge purified EVs were evaluated by DLS. All experiments were repeated at least three times and the representative results are shown.

### Primary keratinocytes obtained from elder subjects represents aged phenotypes even in low passage culture condition

It has been observed that human keratinocytes in primary culture taken from foreskin of children (so-called young keratinocytes in our study) gradually express senescent associated proteins by repeating the passage of cells [15]. To evaluate if the keratinocytes obtained from elder subjects (so-called aged keratinocytes in our study) represent aging phenotypes, we analyzed the expression of several aging marker proteins in young or aged keratinocytes. As shown in Fig. 2A, one of the aging marker proteins, p16, was significantly increased in aged keratinocytes compared to young keratinocytes, even if their passage number were the same (Aged p2 vs Young p2). This induction of p16 was also observed in young keratinocytes but at high passage number (Young p6) which was consistent with the previous report [15]. Moreover, another marker, IDH1, that is known to be reduced during aging process [9], was decreased in aged keratinocytes (Aged p2) and in young keratinocytes (Young p6), but not in young keratinocytes with low passage number (Young p2) (Fig.2C). These data strongly suggest that primary keratinocytes obtained from elder subjects represents aged phenotypes even in low passage number, at least in some aspects.

**Figure 2:**
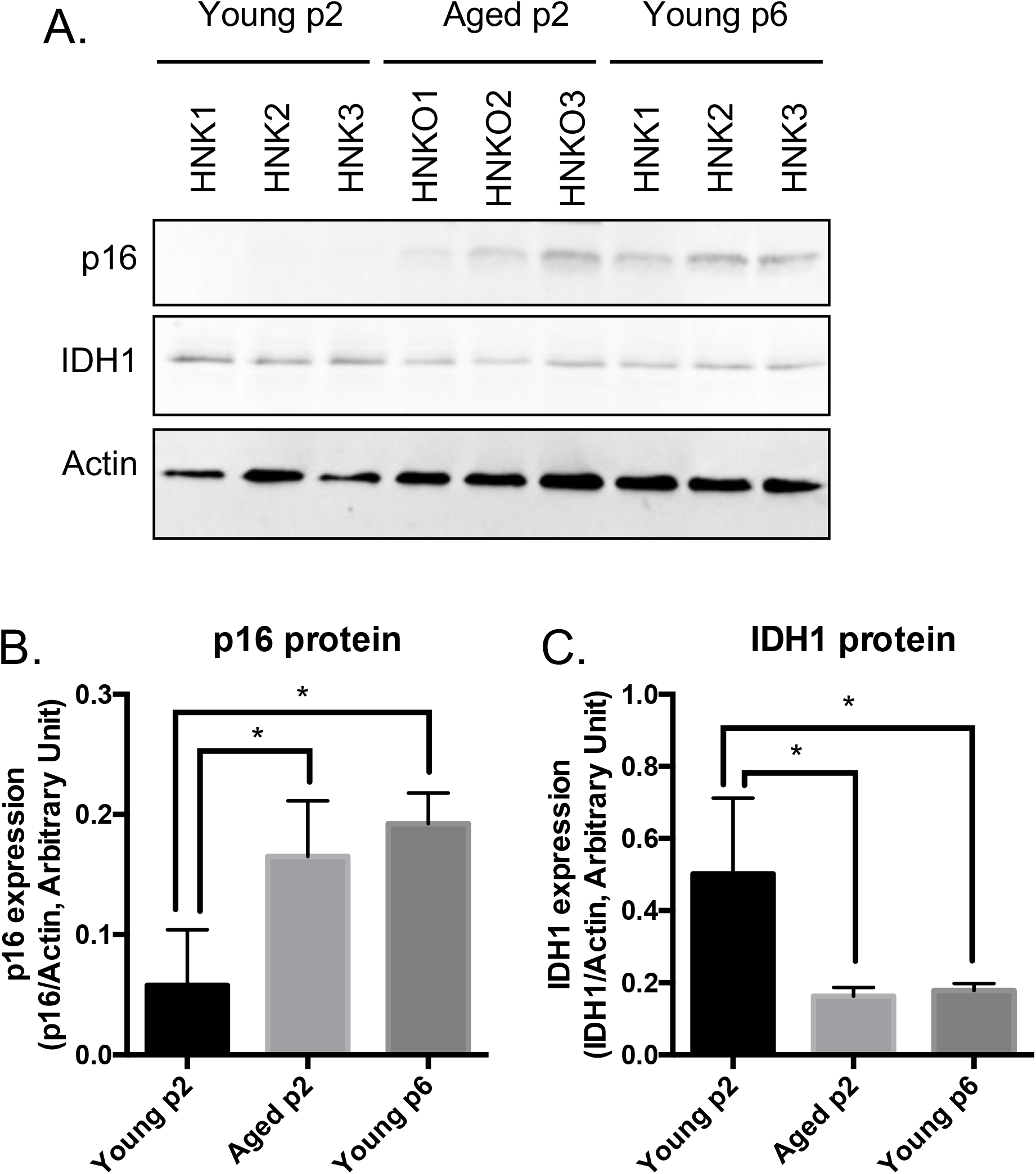
Primary keratinocytes prepared from aged skin represent a valuable model of aging. Primary keratinocytes were isolated from 3 skin biopsies from young (< 5 years) and old individuals (> 60 years). Cells were cultured at early passage (p2) or late passage (p6) and protein were extracted. (A) Western-blot analysis of p16 and IDH1 aging markers. Actin is used as a loading control. (B) Quantification of protein bands from (A). * p-value < 0.05. n=3

### Aged keratinocytes release more EVs compared to young keratinocytes

Then we compared the size and number of small EVs released either from young or aged keratinocytes. Small EV fractions were prepared from cell culture supernatants of either Young p2, Aged p2 or Young p6 keratinocytes, and the obtained vesicles were analyzed by the DLS method as described [16]. The particle size distribution was not significantly different between the three groups (Fig.3A). On the other hand, the relative number of small EVs released from aged keratinocytes (Aged p2), was significantly increased approximately by 2-fold compared to young keratinocytes (Young p2) (Fig.3B). We also measured relative amounts of RNA per vesicle which simply based on the measurement of total RNA amount obtained from each purified fraction, and found that the estimated relative amounts of RNA in vesicles were not significantly changed (Fig.3C). Overall, these results suggested that the number of EVs from keratinocytes were increased upon aging without affecting the size distribution and RNA amounts in vesicles.

**Figure 3:**
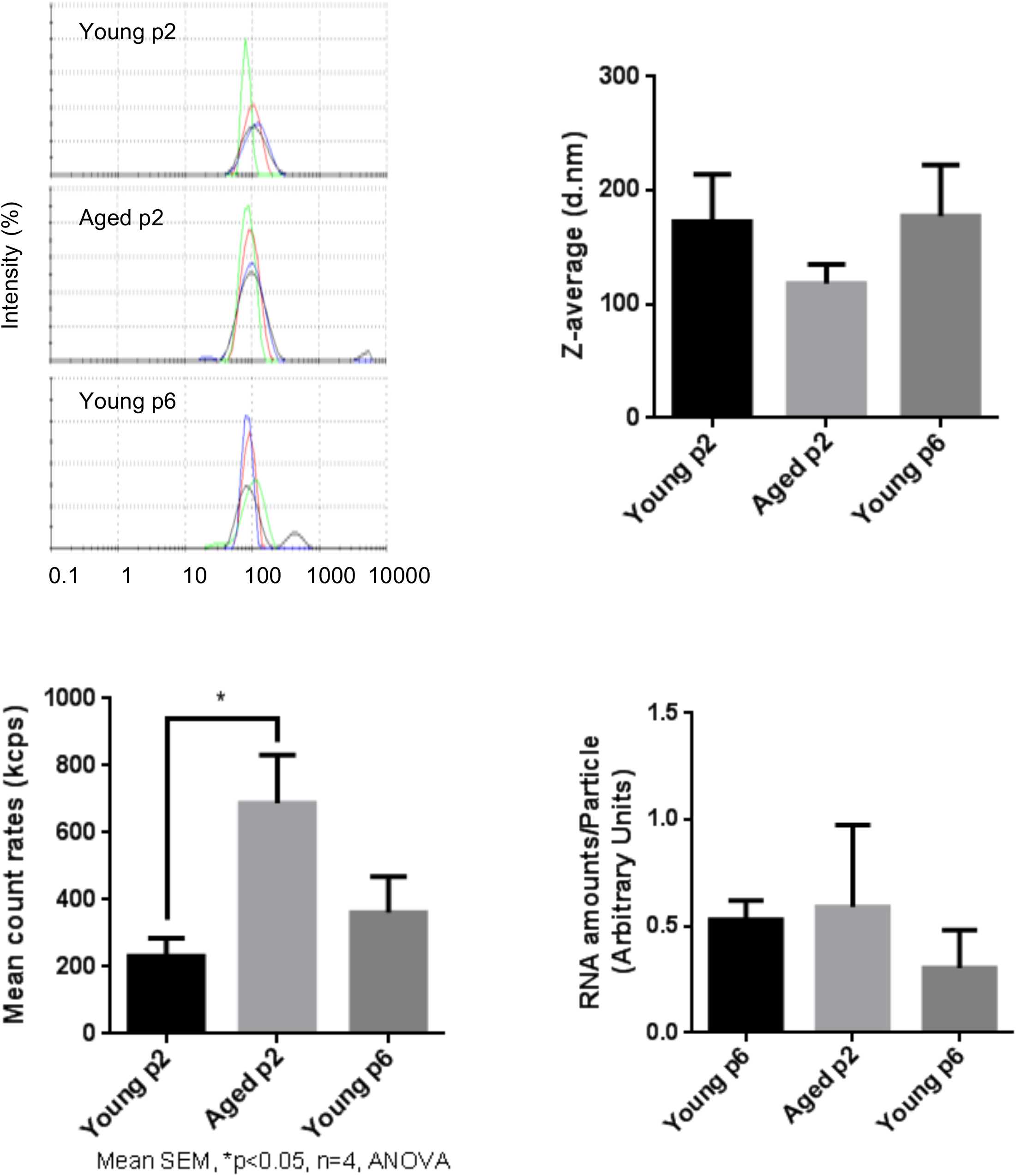
Impact of aging on size and abundance of keratinocytes EVs. EVs were isolated from young p2, old p2 or young p6 keratinocytes culture supernatants. (A) Hydrodynamic size distribution profiles of EVs were evaluated by DLS (right panel). Z-average of EVs were not significantly different between three groups (left panel). (B) Mean count rates of EV samples, with the same attenuator used, were recorded to estimate the abundance of EVs in each fraction. Four independent experiments were performed and the average values ± standard errors are shown in the graph (*p < 0.05, one-way ANOVA with Tukey’s post-hoc test, n=4). (C) Total RNA was isolated from the purified EV fractions and the amounts of RNA were measured as described in Materials and Methods. The amounts of RNA were then divided by the abundance of EVs obtained in (B). Significant difference was not observed between three groups.

### A specific microRNA signature of EVs released from either young or aged-keratinocytes

We have observed that chronological aging impact the abundance of human keratinocytes EVs. The next step was to compare the content of EVs released from young or aged keratinocytes. We focused on microRNAs previously shown to play a key role in EVs function, especially in skin [19, 20].

We next analyzed small RNA profiles in EVs. Total RNA was extracted from independent small EV fractions obtained from young keratinocytes at passage 2 (Young p2), aged keratinocytes at passage 2 (Aged p2), or young keratinocytes at passage 6 (Young p6) (n=4 each) and subjected to small RNA-seq analysis performed by Ion Proton as described in Materials and Methods. We then used several data analysis methods to evaluate miRNA profiles in EVs released from each type of cultured keratinocytes. Unsupervised clustering analysis revealed that three of the four aged samples clustered together (Fig.4A), whereas the number of passages did not account for the observed cluster for the young samples. Principal component analysis was used to identify the more similar samples. Together the first 3 components represent 46% of the variance within the samples. Components 1 (PC1) and 2 (PC2) were able to separate the young p2 from the set of aged p2 and young p6 samples (Fig.4B, left panel). Moreover, the plane formed by the second (PC2) and third components (PC3) can be divided into three zones (young p2, aged p2, young p6 as black, red, blue circle, respectively) (Fig.4B, right panel). Those analysis strongly suggested that the miRNA profiles in EVs could be modulated upon aging process.

**Figure 4:**
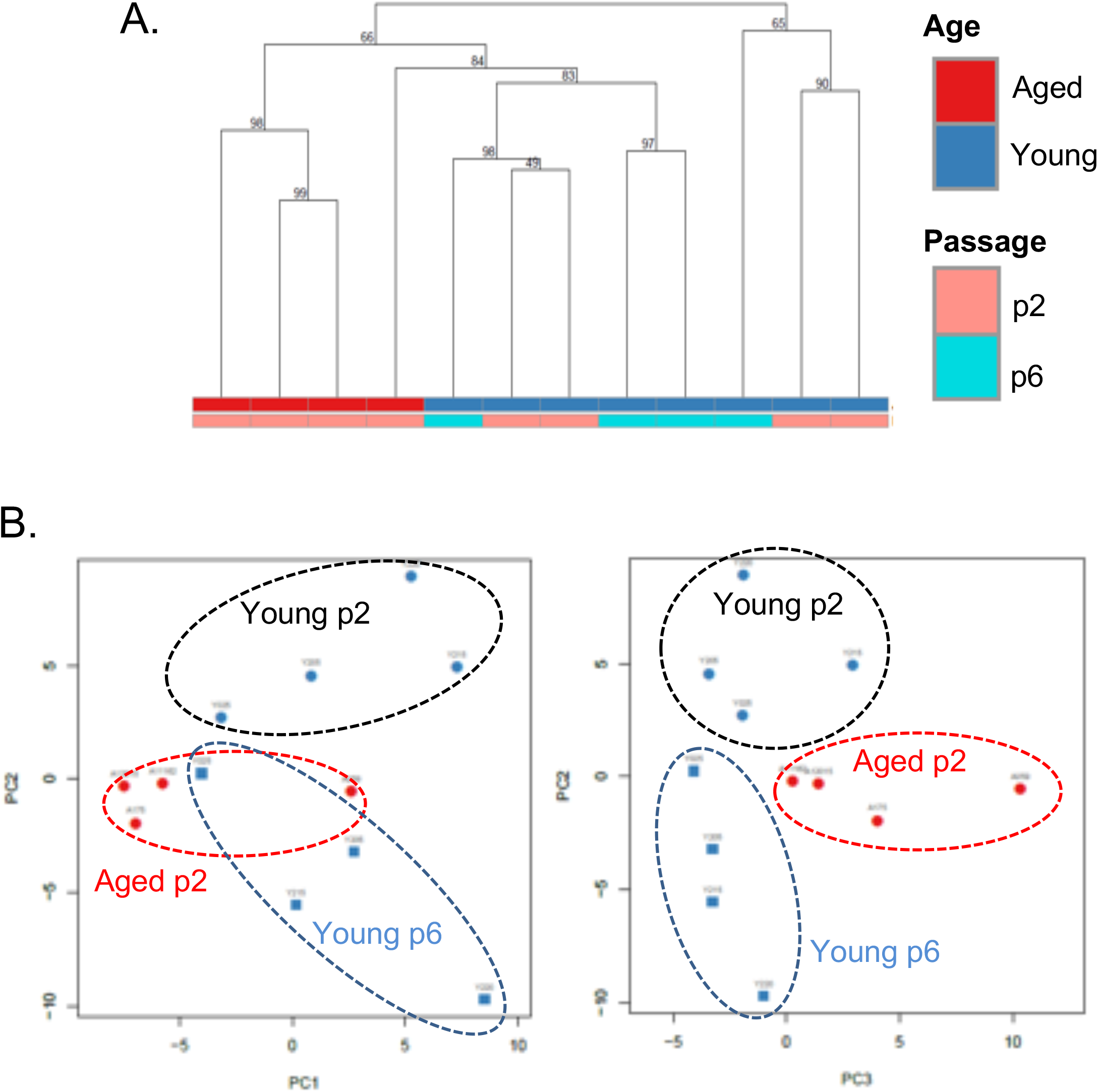
A specific microRNA signature can distinguish EVs from young, aged and senescent keratinocytes. (A) Hierarchical clustering of the EVs samples based on their microRNA signature Keratinocytes supernatants from young or aged skin were used to purify EVs, cultured at early (p2) or late passage (p6). (B) Principal component analysis of the EVs samples based on their microRNA signature, on the left PC1 (19% of the variance) vs PC2 (15 % of the variance), on the right PC2 vs PC3(11% of the variance).

Moreover, miRNA unsupervised analysis revealed EVs miRNAs either upregulated or downregulated during aging and/or senescent process (a heatmap of the 100 miRNAs with highest variance is provided as Supplementary Fig.1). As shown in Table 1, the expression of hsa-miR-30a-5p, miR-30a-3p, and miR-22-3p was increased in aged-p2-EVs, compared to in young-p2-EVs, by approximately 3-fold (adjusted p-value (adj.p) < 0.05, n=4), whereas that of hsa-miR-7977, miR-125a-5p, miR-125b-5p, miR-222-5p, miR-10b-5p, and miR-99b-5p was significantly decreased (adj.p < 0.05, n=4). Partially similar to this data, we also identified EVs miRNAs showing a differential abundance between early (p2) and late (p6) passage young keratinocytes (Table 2): miR-30a-3p and 5p and miR-10-5p were induced in p6 cells whereas miR-125b-3p and 5p, miR-99a-5p, miR-99b-5p and miR-100-5p were repressed. Altogether, these data confirm that chronological aging or replicative senescence modulates the EVs miRNA content.

**Table 1:**
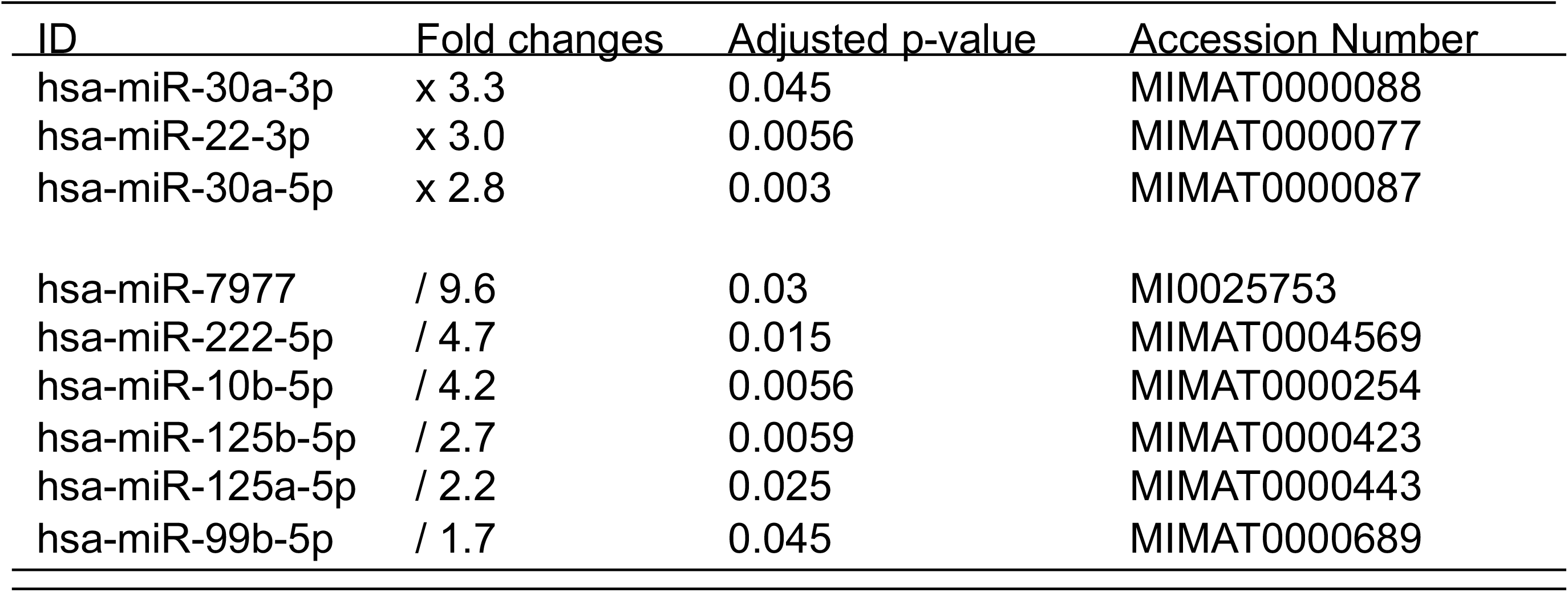
List of microRNAs significantly enriched or depleted in old keratinocytes compared to young keratinocytes at early passage.

| ID | Fold changes | Adjusted p-value | Accession Number |
| --- | --- | --- | --- |
| hsa-miR-30a-3p | x 3.3 | 0.045 | MIMAT0000088 |
| hsa-miR-22-3p | x 3.0 | 0.0056 | MIMAT0000077 |
| hsa-miR-30a-5p | x 2.8 | 0.003 | MIMAT0000087 |
| hsa-miR-7977 | / 9.6 | 0.03 | MI0025753 |
| hsa-miR-222-5p | / 4.7 | 0.015 | MIMAT0004569 |
| hsa-miR-10b-5p | / 4.2 | 0.0056 | MIMAT0000254 |
| hsa-miR-125b-5p | / 2.7 | 0.0059 | MIMAT0000423 |
| hsa-miR-125a-5p | / 2.2 | 0.025 | MIMAT0000443 |
| hsa-miR-99b-5p | / 1.7 | 0.045 | MIMAT0000689 |

**Table 2:**
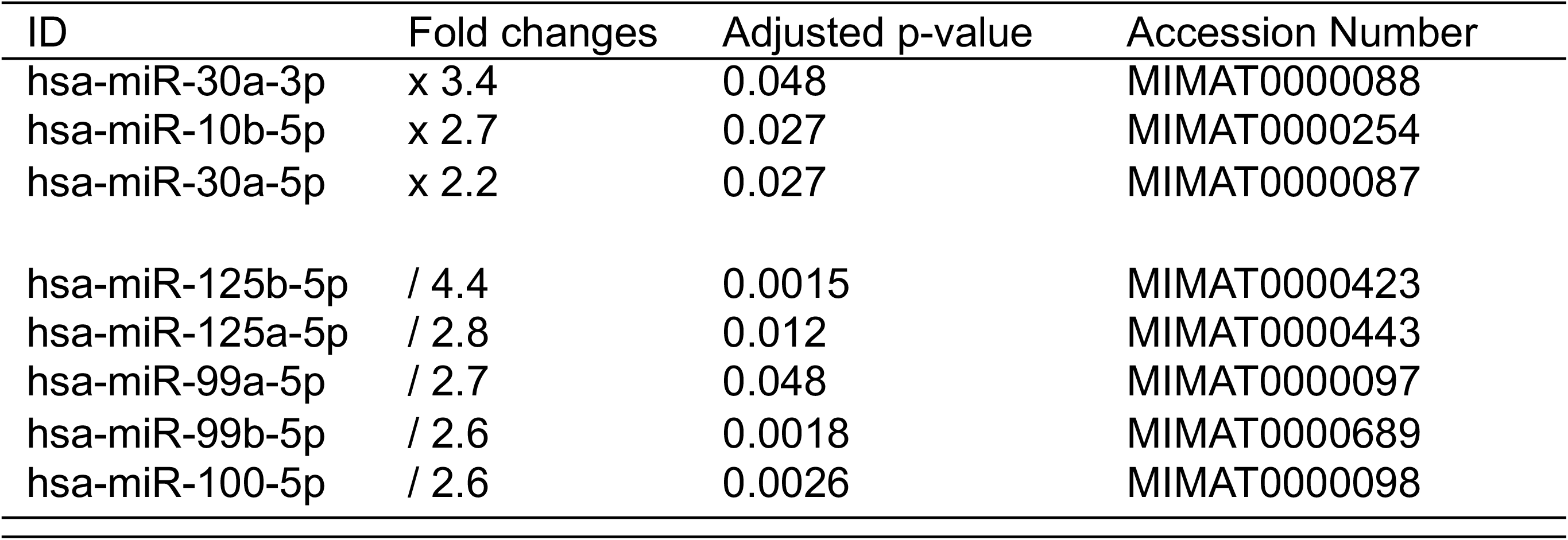
List of miRNAs significantly enriched or depleted in young keratinocytes at early passage compared to young keratinocytes at late passage.

### Aged keratinocytes-derived EVs impact the proliferation and migration of young keratinocytes

In order to elucidate the role of EVs released from keratinocytes, we first confirmed whether purified EVs had the potential to fuse with other cultured keratinocytes. Purified EVs were florescent labeled with CM-Dil, then added on keratinocyte culture for 24h. Cells were then washed with PBS to remove non-fused EVs and observed by fluorescent microscopy. Even though the fluorescent intensity was variable among cells, fluorescent positive keratinocytes were observed when adding CM-Dil-labeled EVs, which suggested that purified EVs released from keratinocytes fused with keratinocytes themselves (Fig.5A). We then analyzed the effect of EVs treatment on young keratinocytes proliferation. Cultured keratinocytes at early passage and plated at low density were treated with small EVs from young or aged keratinocytes. Their proliferation rate was assessed by DNA content measurement 24h to 72h after the treatment. A significant reduction of keratinocytes proliferation was observed at 48h and 72h in the cells treated with old EVs whereas the treatment with young EVs had no measurable effect (Fig.5B). Next, to go further into the possible effects of old EVs on young keratinocytes, we measured by quantitative RT-PCR the transcript expression of several genes related to cell proliferation, differentiation and aging process. As shown in Fig.5C, the treatment with EVs significantly induced the gene expression of the anti-proliferative regulator p21 and might activate the senescence marker p16 whereas the checkpoint regulator p53 was not modulated. Consistent with a decreased of cell proliferation, the early differentiation markers K1 and K10 were induced as well as the late differentiation marker loricrin. This suggest that the young keratinocytes treated with old EVs enter into a transition between proliferation and differentiation. We next checked whether the old EVs treatment affected young keratinocytes migration in a scratch wound-healing assay. As shown on Fig.6, a strong reduction of the migration speed was observed for keratinocytes treated with old EVs: at 40h the wound was completely closed in the control condition whereas in the treated condition, the scratch was still largely opened.

**Figure 5:**
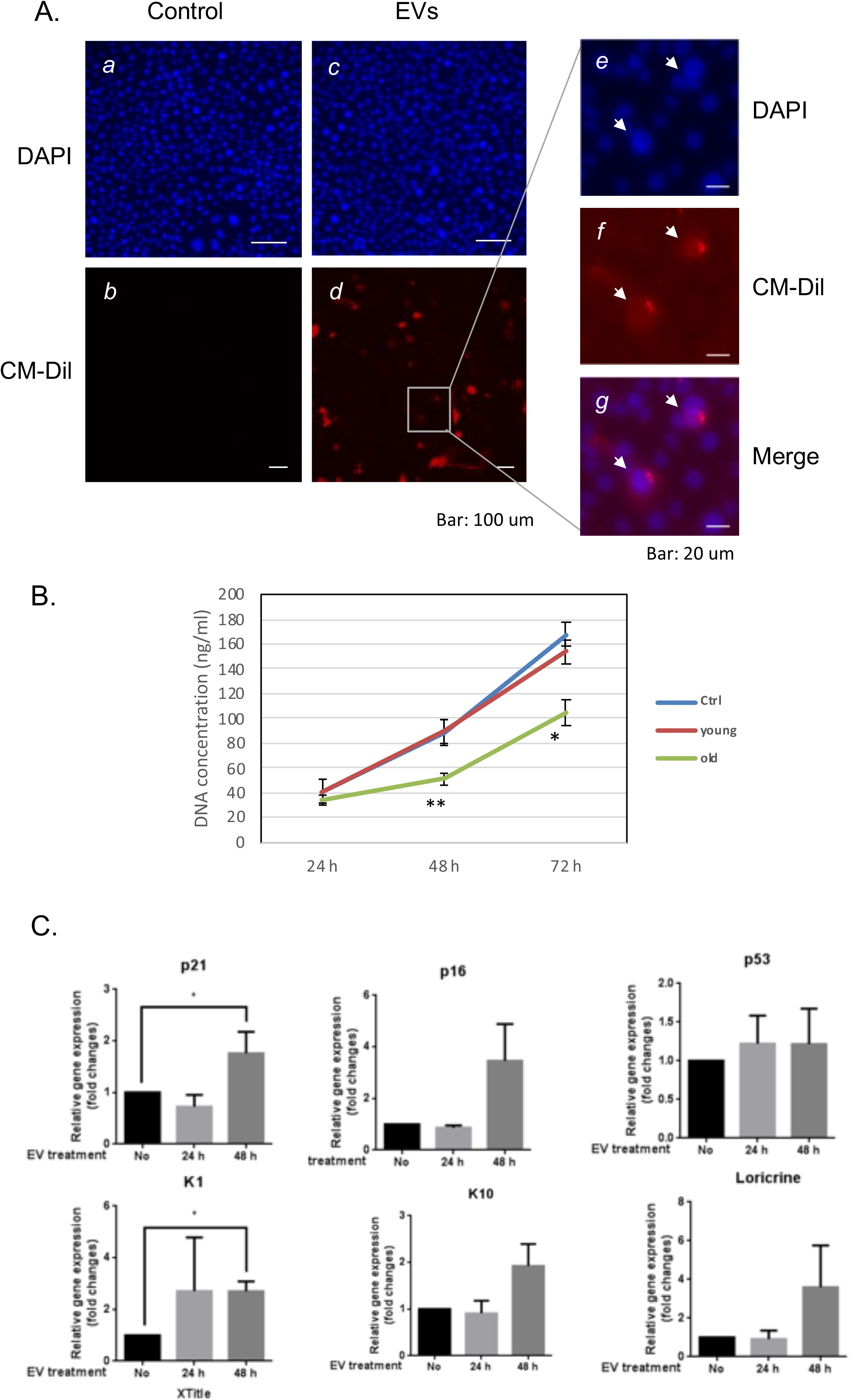
Aged EVs impact proliferation of young keratinocytes in 2D culture. (A) Primary keratinocytes were treated with DAPI only (a, c) or by DAPI plus fluorescently labeled (CM-Dil) EVs from young keratinocytes culture medium (b, d). A part of image d is shown with higher magnification in e, f and g. (B) Proliferation profile of young cultured keratinocytes treated with EVs purified from young or aged keratinocytes. DNA concentration in treated keratinocytes was measured 24h, 48h and 72h after the beginning of the treatment. * p-value < 0.05. ** p-value < 0.01. n=3 (C) Expression analysis of transcripts for P21, P16, P53, KRT1 (K1), KRT10 (K10) and loricrin in young keratinocytes treated with EVs from old keratinocytes. Data were normalized to the non-treated condition. * p-value < 0.05. n=3.

**Figure 6:**
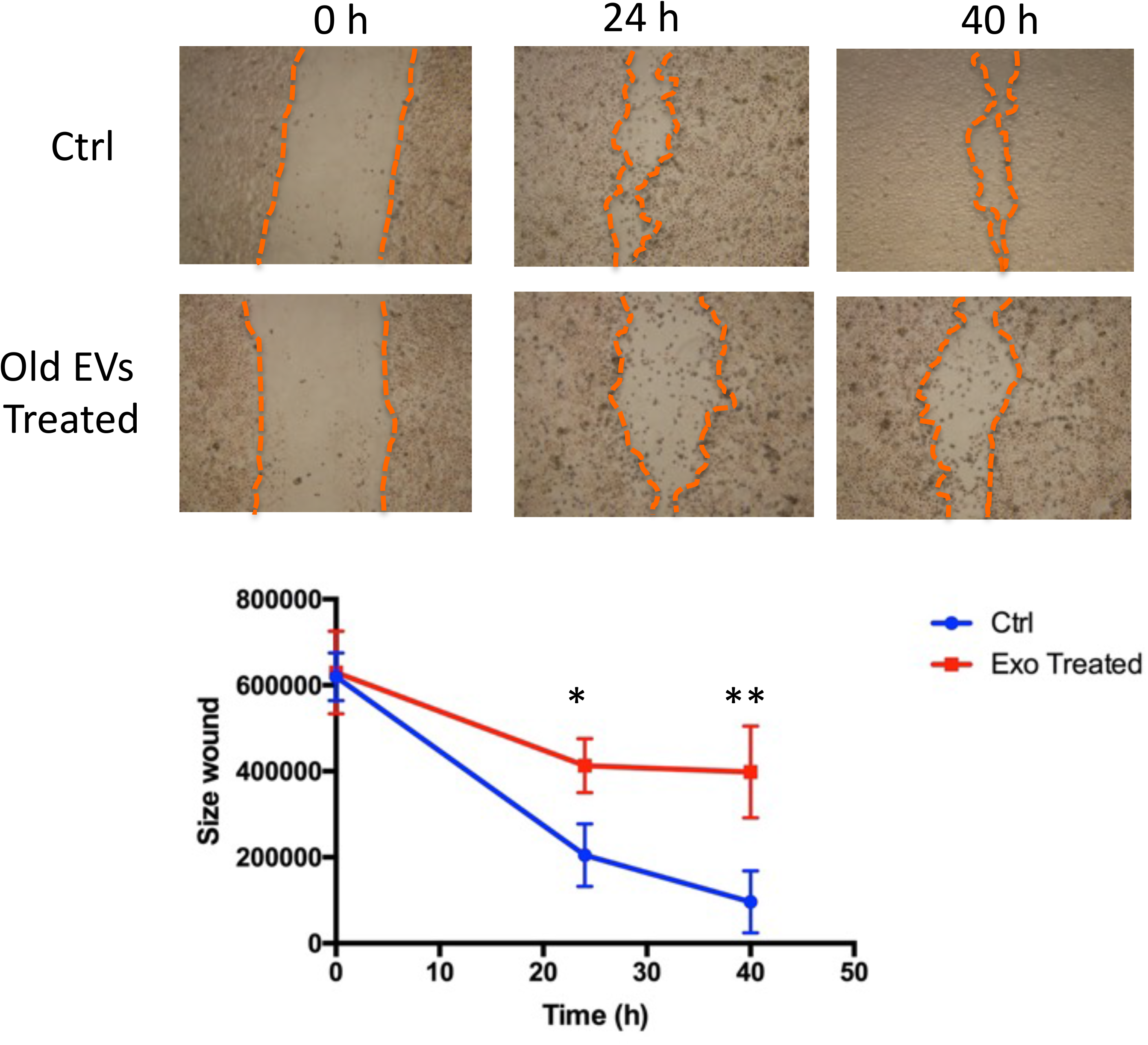
Aged EVs impact migration of young keratinocytes in 2D culture. Confluent young keratinocytes cultures were scratch wounded and exposed to old keratinocytes EVs for 40h. The remaining wound area were quantified at 24h and 40h by ImageJ from triplicate experiments (* p-value < 0.05. ** p-value < 0.01. Student’s t-test).

Overall, these results indicate that small EVs derived from human keratinocytes are still functional and are able to impair keratinocytes proliferation and migration in a monolayer culture system.

### Old keratinocytes-derived EVs impact epidermal reconstruction in an organotypic 3D model

We then investigated the effect of EVs treatment on the keratinocyte ability to reconstruct an epidermal tissue *in vitro*. We first used keratinocytes from female abdominal skin (>60 years). After an initial 48h immerged culture allowing keratinocytes proliferation, the epidermis was placed at the air-liquid interface for 14 days to permit keratinocytes differentiation. Since the first days after seeding cells, we treated the tissue with EVs collected from young keratinocytes conditioned medium. The opposite experiment was also done: using young keratinocytes extracted from infant foreskin and EVs from aged keratinocytes conditioned medium. The histological aspects of these tissues are presented on Fig.7A. We first observed that the reconstructed human epidermis (RHE) obtained from aged keratinocytes were thinner than those obtained from young keratinocytes (Fig.7A), indicating that the proliferation-differential potential of aged keratinocytes was reduced compared to young cells. We then studied the impact of EVs treatments. For RHE obtained from old cells treated with young EVs, no difference in term of thickness were visible. This observation was confirmed by the measurement of epidermal thickness (Fig.7B), with no significant difference between the two conditions. On the contrary, in the reverse experiment, we observed a thinner tissue (Fig.7A) for the epidermis treated with aged EVs. The epidermal thickness was also measured and we were able to confirm a significant decrease of the tissue thickness (Fig.7C) (t.test, p<0,0001).

**Figure 7:**
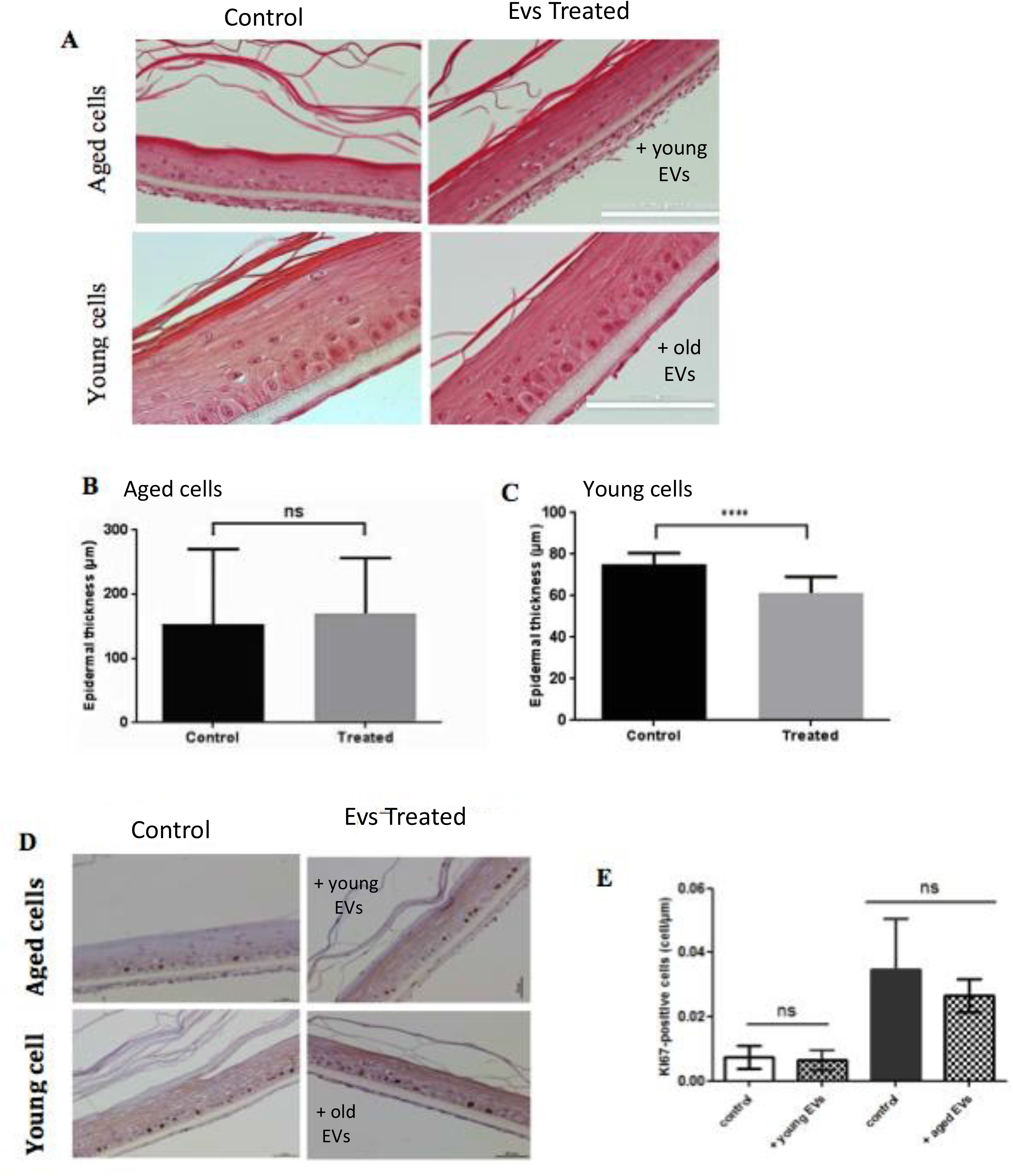
Aged EVs impact epidermal reconstruction in a 3D model. Young or old keratinocytes were used to reconstruct cultured epidermis in a 3D organotypic model. Reconstructed epidermis obtained with aged keratinocytes were treated with EVs obtained from young keratinocytes culture medium. Conversely, reconstructed epidermis obtained with young keratinocytes were treated with EVs obtained from old keratinocytes culture medium. Representative sections stained with hematoxylin/eosin/safranin are shown in A (Scale bar = 100 μm). The epidermal thickness in 3D culture corresponding to old keratinocytes treated with young EVs are shown in B (ns: non-significant in Student t-test). The epidermal thickness in 3D culture corresponding to young keratinocytes treated with old EVs are shown in C (*** p-value < 0.001. Student T-test). The reconstructed epidermis sections were stained with Ki67. Representative images of the various conditions are shown in D. The number of Ki-67 positive cells per μm is shown in E (ns: non-significant in a Student t-test).

These effect on epidermis thickness might be due to a difference in term of proliferation within the different conditions. To assess this point, we investigated the expression of the Ki67 proliferation marker in our RHE sections. As expected, we observed proliferative cells only in the basal layer of the epidermis (Fig.7D) and measured an increased number of proliferative cells in the RHE prepared with young keratinocytes (Fig.6E right bar). However, we didn’t observe any significant effect of EVs treatment on Ki67 expression after 14 days of differentiation. This might suggest a possible earlier effect in the first steps of keratinocytes proliferation during the emerged phase of the epidermal reconstruction. To decipher a possible effect of old EVs treatment on epidermal differentiation, we analyzed the expression of the differentiation markers KRT1 and involucrin in the RHEs. For the RHE obtained from young keratinocytes treated with aged EVs, we observed a thinner epidermis with a reduced surface of staining for the 2 markers but no significant difference in term of labeling intensity (Supplementary Fig.2).

### Old keratinocytes-derived EVs slow down early phases of wound healing in mice

To decipher the *in vivo* impact of old EVs on keratinocyte migration and proliferation, we explored their influence on wound healing, a multi-step process involving keratinocytes migration and proliferation in the early phase of re-epithelization. We used a protocol already developed to study the impact of human EVs from various cell types on wound healing in mice [17, 18]. EVs purified from old keratinocytes were injected around the excision wound and the progress of the wound closure was then followed by image analysis during the 12 days after injection. In the group of mice treated by injection of old keratinocytes-derived EVs, we observed a strong slow-down of the wound closure in the first days compared to controls injected only with PBS (Fig.8A), statistically significant for day 1 to day 4 (Fig.8B). After 10 days, the percentage of wound closure was quite similar in the two conditions (Fig.8B). Those results indicate that human keratinocytes EVs are able to modulate wound healing in mice and suggest that old keratinocytes EVs influence the early phases of re-epithelization, characterized by proliferation and migration of the keratinocytes from the wound edge.

**Figure 8:**
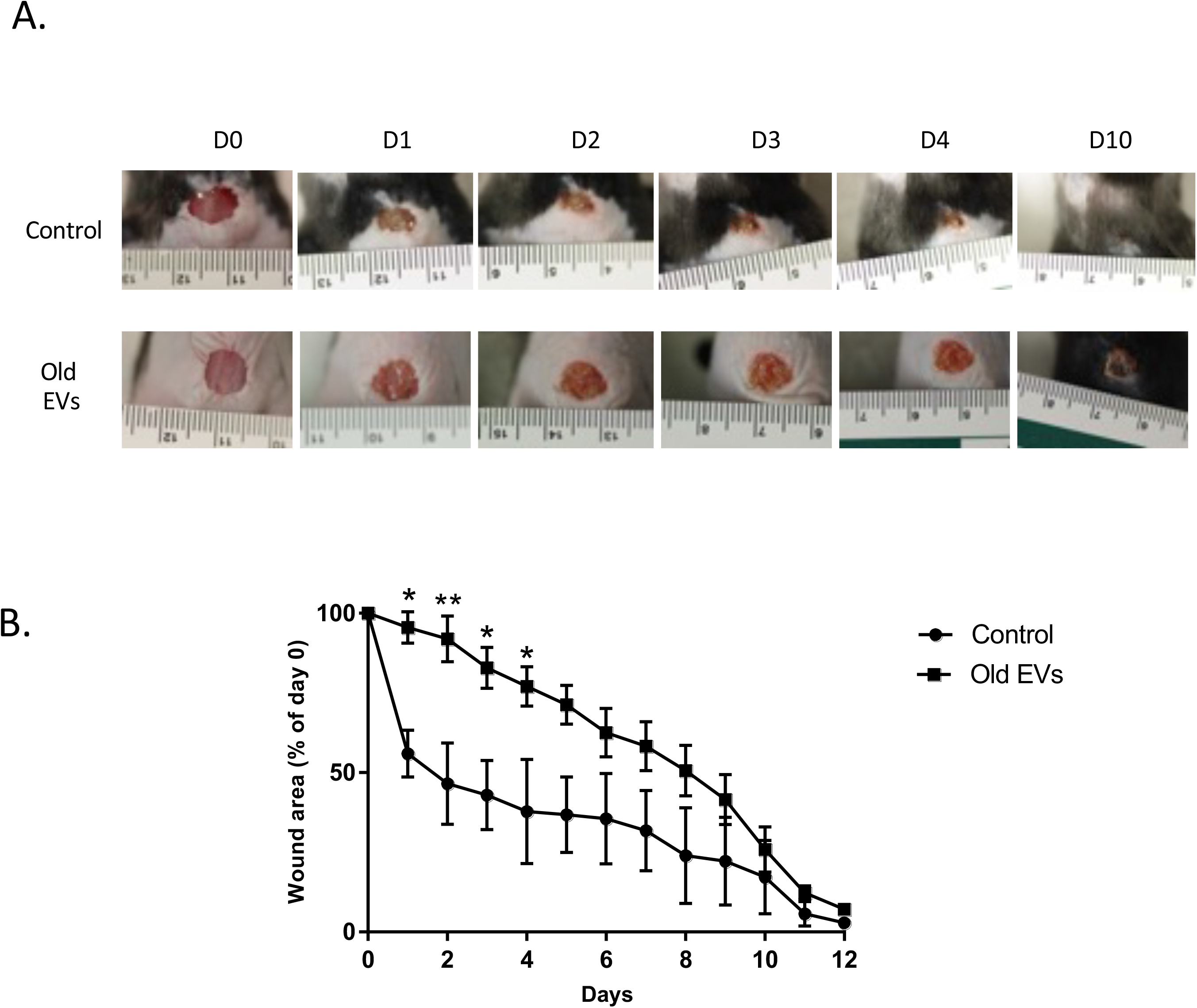
Aged EVs impact the early phases of wound healing in mice. Wound closure assay on mice treated with EVs from old keratinocytes. (A) Representative photographs of full-thickness excisional wounds treated at day 0 (D0) with NaCl (Control n=4) or with EVs from old human keratinocytes (n=8) every day from day 0 to day 10 (D10). (B) Quantitative analysis of wound healing closure in each group, expressed as a percentage of the initial wound area (* p-value < 0.05. ** p-value < 0.01. Student’s t-test).

## Discussion

In this article, we describe for the first time the impact of chronological aging on EVs production by human keratinocytes. Keratinocytes are the constitutive cells of the epidermis, which ensures the barrier function of the skin which is impaired in aged skin [21]. It is then necessary to better understand how chronological aging shapes keratinocytes function, especially how these cells ensure epidermis renewal and differentiation. We have shown here that cultured primary keratinocytes from young or aged skin represent a valuable model to study the impact of aging. Indeed, keratinocytes from old skin express aging markers such as p16 and show a lower ability to reconstruct an epidermis in a 3D model. Even in culture in a rich medium, keratinocytes from old skin retains some features of the tissue they are extracted from and can be used to decipher the impact of aging at the cellular level.

Concerning EVs production by epidermal keratinocytes, it have been shown that exosomes released by keratinocytes are able to modulate genes expression in fibroblasts [22] and to control melanin production by melanocytes [23] but the effect of keratinocytes EVs on keratinocytes themselves has never been investigated. We confirmed here that keratinocytes EVs can fused and be integrated by keratinocytes themselves and we observed that EVs from aged keratinocytes (i.e. keratinocytes extracted from aged individual skin) are able to modulate the proliferation and the migration of young keratinocytes in monolayer culture. Moreover, we observed that EVs secreted from old keratinocytes were also able to modulate epidermis reconstruction in an organotypic 3D model and to influence the early phases of wound healing in mice. These data demonstrate that the EVs produced by aged cells exert specific effects, different to those produced by young cells. This also suggests that the keratinocytes EV content is modified in aged cells. Here we focused on miRNAs that represent important mediators of EVs functional effect, especially in aging [24]. We observed a specific EV miRNAs signature in aged compared to young keratinocytes and identified several miRNAs significantly enriched or depleted in EVs from old cells. This was the case of miR-30a, for which the two strands −3p and −5p were enriched in aged keratinocytes EVs. We previously identified miR-30a as a key regulator of human epidermal aging: miR-30a is actually up-regulated in aged epidermis and this induction is at least partly responsible for age-related differentiation defects and cell death boost in human epidermis [9]. The fact that miR-30a-3p and 5p are carried by EVs and enriched in EVs from old cells suggests that aged keratinocytes could propagate their functional defects to surrounding cells by miR-30a secretion. However, miR-30a might not be involved in the anti-proliferative effect observed in old-EVs treated cells; since in our previous study we didn’t observe any modification of the proliferation status in miR-30a over-expressing cells [9]. MiR-22-3p is another miRNA that we found enriched in old keratinocytes EVs. This microRNA, already shown induced by aging in vascular endothelial cells [25], is a potential negative regulator of proliferation and migration as observed in various normal or cancer cell types [26–28]. The real functional involvement of miR-22-3p in the aged EVs effect needs to be clarified by further specific experiments but this is a good candidate to regulate proliferation and migration in human keratinocytes. Among the 6 microRNAs found significantly depleted in exosomes from old cells, we were particularly interested by miR-125b-5p because this microRNA has been already found repressed by chronological aging in skin cells [29] and acts as a potential marker of epidermal stem cells and regulator of their proliferation potential [30]. The miR-125b depleted EVs might therefore exerted a negative effect on the progenitor population of keratinocytes by modulating their proliferation, similarly to a previous report in dermal fibroblasts [31].

We also observed that the EVs abundance was increased in aged cells. To our knowledge this has been not clearly observed so far for chronological aging. Several groups have observed an increase of the EVs release in senescent cells [32, 33], EVs being part of the secretory activity of these cells also called SASP for Senescent Associated Secretory Phenotype [34]. However, senescence is only one of the numerous mechanisms explaining the aging changes and the effect of the chronological aging on EV production need to be studied in its entirety. In few studies, the effect of aging on EVs size and abundance has been evaluated: Davis et al. didn’t observe any changes in mouse bone marrow EVs concentration and size with aging, whereas their miRNAs content was altered [35]. Alibhai et al. observed an elevated level of plasma EVs in old mice [36]. On the contrary, the abundance of circulating plasma EVs was decreased with advancing age in human samples but the EVs size distribution was not impacted [37]. These contradictory observations suggest that the effect of aging on EVs secretion is mainly tissuespecific.

In this study we characterized the miRNA content of EVs secreted by various types of cultured human keratinocytes: keratinocytes from young skin at early and late passage, keratinocytes from old skin at early passage. We were able to compare the impact of replicative senescence and chronological aging on the abundance and microRNAs content of secreted EVs. As shown in Fig.4A, the global microRNA signature of the EVs from aged cells is clearly different to the one from young cells whatever their number of passages might be, suggesting that replicative senescence do not fully mimic the phenotype of chronological aging in keratinocytes. However, when we consider the microRNAs that are significantly modulated between old/young or early/late passages, several microRNAs, including miR-30a-3p and 5p, miR-99b-5p, miR-125a and b are modulated in the same way by chronological aging and by replicative senescence as well. This suggests that the differential loading of these microRNAs in keratinocytes EVs could be linked to molecular pathways in common between senescence cells and aged cells.

Another interesting question is whether the abundance of the microRNAs in the EVs reflect the total abundance of the microRNAs in the secreting cells. In other words, is the EVs miRnome a mirror of the cellular miRnome? We were able to analyze the microRNA content of 2 samples of young keratinocytes at early passage (data not shown). A rapid comparison between the cellular and EVs microRNA signatures reveals strong differences in term of microRNAs relative abundance in accordance with previous observations showing that the microRNAs selection and incorporation in EVs is an active and selective mechanism [38, 39].

To conclude, we have shown here that aging modulates EVs abundance, function and microRNA content in human keratinocytes. We have observed that EVs from aged keratinocytes exert an anti-proliferative and anti-migratory effect on young keratinocytes, and restraint their ability to reconstruct an epidermis in a 3D model. Moreover, EVs from aged keratinocytes delays wound healing in mice. EVs from old cells might be then deleterious for their surrounding cells in the tissue. EVs could then be a part of a more global secretory phenotype able to negatively modulate epidermis function in aged skin. Further studies will be necessary to fully characterize this message secreted by aged cells. Conversely, we didn’t observe any beneficial effect of EVs from young cells on keratinocytes proliferation and epidermal reconstruction. Contrary to what have been observed for EVs produced by different types of skin resident stem cells, such as mesenchymal [40] or adipocytes stem cells [41], EVs from young keratinocytes are not able to reverse the deleterious effect of aging in epidermis and definitely do not represent valuable biological compounds for anti-aging therapeutic strategies.

---

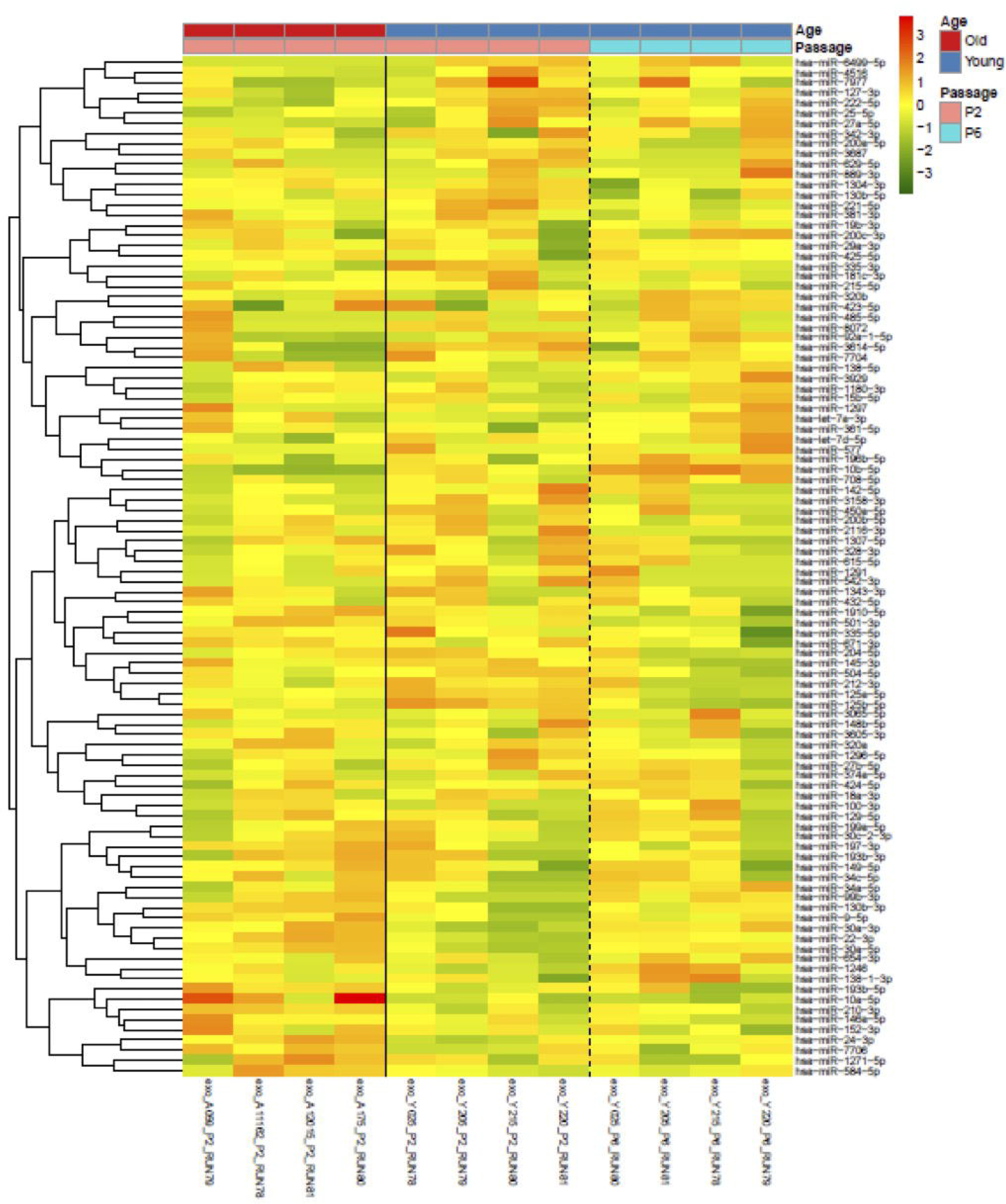

## Notes

### Competing Interest Statement

The authors have declared no competing interest.

## References

1. Gilhar A, Ullmann Y, Karry R, Shalaginov R, Assy B, Serafimovich S, Kalish RS. Ageing of human epidermis: the role of apoptosis, Fas and telomerase. Br J Dermatol. 2004; 150: 56–63.

2. Rinnerthaler M, Duschl J, Steinbacher P, Salzmann M, Bischof J, Schuller M, Wimmer H, Peer T, Bauer JW, Richter K. Age-related changes in the composition of the cornified envelope in human skin. Exp Dermatol. 2013; 22: 329–35.

3. Lopez-Otin C, Blasco MA, Partridge L, Serrano M, Kroemer G. The hallmarks of aging. Cell. 2013; 153: 1194–217.

4. van Deursen JM. The role of senescent cells in ageing. Nature. 2014; 509: 439–46.

5. Nelson G, Wordsworth J, Wang C, Jurk D, Lawless C, Martin-Ruiz C, von Zglinicki T. A senescent cell bystander effect: senescence-induced senescence. Aging Cell. 2012; 11: 345–9.

6. Waldera Lupa DM, Kalfalah F, Safferling K, Boukamp P, Poschmann G, Volpi E, Gotz-Rosch C, Bernerd F, Haag L, Huebenthal U, Fritsche E, Boege F, Grabe N, et al. Characterization of Skin Aging-Associated Secreted Proteins (SAASP) Produced by Dermal Fibroblasts Isolated from Intrinsically Aged Human Skin. J Invest Dermatol. 2015; 135: 1954–68.

7. McBride JD, Rodriguez-Menocal L, Badiavas EV. Extracellular Vesicles as Biomarkers and Therapeutics in Dermatology: A Focus on Exosomes. J Invest Dermatol. 2017; 137: 1622–9.

8. Stahl PD, Raposo G. Extracellular Vesicles: Exosomes and Microvesicles, Integrators of Homeostasis. Physiology (Bethesda). 2019; 34: 169–77.

9. Muther C, Jobeili L, Garion M, Heraud S, Thepot A, Damour O, Lamartine J. An expression screen for aged-dependent microRNAs identifies miR-30a as a key regulator of aging features in human epidermis. Aging (Albany NY). 2017; 9: 2376–96.

10. Franco N, Lamartine J, Frouin V, Le Minter P, Petat C, Leplat JJ, Libert F, Gidrol X, Martin MT. Low-dose exposure to gamma rays induces specific gene regulations in normal human keratinocytes. Radiat Res. 2005; 163: 623–35.

11. Lasser C, Eldh M, Lotvall J. Isolation and characterization of RNA-containing exosomes. J Vis Exp. 2012: e3037.

12. Baras AS, Mitchell CJ, Myers JR, Gupta S, Weng LC, Ashton JM, Cornish TC, Pandey A, Halushka MK. miRge - A Multiplexed Method of Processing Small RNA-Seq Data to Determine MicroRNA Entropy. PLoS One. 2015; 10: e0143066.

13. Wu Y, Deng W, Klinke DJ, 2nd. Exosomes: improved methods to characterize their morphology, RNA content, and surface protein biomarkers. Analyst. 2015; 140: 6631–42.

14. Konoshenko MY, Lekchnov EA, Vlassov AV, Laktionov PP. Isolation of Extracellular Vesicles: General Methodologies and Latest Trends. Biomed Res Int. 2018; 2018: 8545347.

15. Gosselin K, Deruy E, Martien S, Vercamer C, Bouali F, Dujardin T, Slomianny C, Houel-Renault L, Chelli F, De Launoit Y, Abbadie C. Senescent keratinocytes die by autophagic programmed cell death. Am J Pathol. 2009; 174: 423–35.

16. van der Pol E, Hoekstra AG, Sturk A, Otto C, van Leeuwen TG, Nieuwland R. Optical and non-optical methods for detection and characterization of microparticles and exosomes. J Thromb Haemost. 2010; 8: 2596–607.

17. Zhao B, Zhang Y, Han S, Zhang W, Zhou Q, Guan H, Liu J, Shi J, Su L, Hu D. Exosomes derived from human amniotic epithelial cells accelerate wound healing and inhibit scar formation. J Mol Histol. 2017; 48: 121–32.

18. Hu L, Wang J, Zhou X, Xiong Z, Zhao J, Yu R, Huang F, Zhang H, Chen L. Exosomes derived from human adipose mensenchymal stem cells accelerates cutaneous wound healing via optimizing the characteristics of fibroblasts. Sci Rep. 2016; 6: 32993.

19. Yan H, Gao Y, Ding Q, Liu J, Li Y, Jin M, Xu H, Ma S, Wang X, Zeng W, Chen Y. Exosomal Micro RNAs Derived from Dermal Papilla Cells Mediate Hair Follicle Stem Cell Proliferation and Differentiation. Int J Biol Sci. 2019; 15: 1368–82.

20. Liu Y, Xue L, Gao H, Chang L, Yu X, Zhu Z, He X, Geng J, Dong Y, Li H, Zhang L, Wang H. Exosomal miRNA derived from keratinocytes regulates pigmentation in melanocytes. J Dermatol Sci. 2019; 93: 159–67.

21. Luebberding S, Krueger N, Kerscher M. Age-related changes in skin barrier function - quantitative evaluation of 150 female subjects. Int J Cosmet Sci. 2013; 35: 183–90.

22. Huang P, Bi J, Owen GR, Chen W, Rokka A, Koivisto L, Heino J, Hakkinen L, Larjava H. Keratinocyte Microvesicles Regulate the Expression of Multiple Genes in Dermal Fibroblasts. J Invest Dermatol. 2015; 135: 3051–9.

23. Lo Cicero A, Delevoye C, Gilles-Marsens F, Loew D, Dingli F, Guere C, Andre N, Vie K, van Niel G, Raposo G. Exosomes released by keratinocytes modulate melanocyte pigmentation. Nat Commun. 2015; 6: 7506.

24. Xu D, Tahara H. The role of exosomes and microRNAs in senescence and aging. Adv Drug Deliv Rev. 2013; 65: 368–75.

25. Takeda E, Suzuki Y, Sato Y. Age-associated downregulation of vasohibin-1 in vascular endothelial cells. Aging Cell. 2016; 15: 885–92.

26. Huang SC, Wang M, Wu WB, Wang R, Cui J, Li W, Li ZL, Li W, Wang SM. Mir-22-3p Inhibits Arterial Smooth Muscle Cell Proliferation and Migration and Neointimal Hyperplasia by Targeting HMGB1 in Arteriosclerosis Obliterans. Cell Physiol Biochem. 2017; 42: 2492–506.

27. Liu Y, Li H, Liu Y, Zhu Z. MiR-22-3p targeting alpha-enolase 1 regulates the proliferation of retinoblastoma cells. Biomed Pharmacother. 2018; 105: 805–12.

28. Zhao L, Wang Y, Liu Q. Catalpol inhibits cell proliferation, invasion and migration through regulating miR-22-3p/MTA3 signalling in hepatocellular carcinoma. Exp Mol Pathol. 2019; 109: 51–60.

29. Toyokuni S, Jiang L, Wang S, Hirao A, Wada T, Soh C, Toyama K, Kawada A. Aging rather than sun exposure is a major determining factor for the density of miR-125b-positive epidermal stem cells in human skin. Pathol Int. 2015; 65: 415–9.

30. Zhang L, Stokes N, Polak L, Fuchs E. Specific microRNAs are preferentially expressed by skin stem cells to balance self-renewal and early lineage commitment. Cell Stem Cell. 2011; 8: 294–308.

31. Kozlova A, Pachera E, Maurer B, Jungel A, Distler JHW, Kania G, Distler O. MicroRNA-125b Regulates Fibroblast Apoptosis Proliferation in Systemic Sclerosis. Arthritis Rheumatol. 2019.

32. Lehmann BD, Paine MS, Brooks AM, McCubrey JA, Renegar RH, Wang R, Terrian DM. Senescence-associated exosome release from human prostate cancer cells. Cancer Res. 2008; 68: 7864–71.

33. Takasugi M, Okada R, Takahashi A, Virya Chen D, Watanabe S, Hara E. Small extracellular vesicles secreted from senescent cells promote cancer cell proliferation through EphA2. Nat Commun. 2017; 8: 15729.

34. Kadota T, Fujita Y, Yoshioka Y, Araya J, Kuwano K, Ochiya T. Emerging role of extracellular vesicles as a senescence-associated secretory phenotype: Insights into the pathophysiology of lung diseases. Mol Aspects Med. 2018; 60: 92–103.

35. Davis C, Dukes A, Drewry M, Helwa I, Johnson MH, Isales CM, Hill WD, Liu Y, Shi X, Fulzele S, Hamrick MW. MicroRNA-183-5p Increases with Age in Bone-Derived Extracellular Vesicles, Suppresses Bone Marrow Stromal (Stem) Cell Proliferation, and Induces Stem Cell Senescence. Tissue Eng Part A. 2017; 23: 1231–40.

36. Alibhai FJ, Lim F, Yeganeh A, DiStefano PV, Binesh-Marvasti T, Belfiore A, Wlodarek L, Gustafson D, Millar S, Li SH, Weisel RD, Fish JE, Li RK. Cellular senescence contributes to age-dependent changes in circulating extracellular vesicle cargo and function. Aging Cell. 2020; 19: e13103.

37. Eitan E, Green J, Bodogai M, Mode NA, Baek R, Jorgensen MM, Freeman DW, Witwer KW, Zonderman AB, Biragyn A, Mattson MP, Noren Hooten N, Evans MK. Age-Related Changes in Plasma Extracellular Vesicle Characteristics and Internalization by Leukocytes. Sci Rep. 2017; 7: 1342.

38. Janas T, Janas MM, Sapon K, Janas T. Mechanisms of RNA loading into exosomes. FEBS Lett. 2015; 589: 1391–8.

39. Li C, Qin F, Hu F, Xu H, Sun G, Han G, Wang T, Guo M. Characterization and selective incorporation of small non-coding RNAs in non-small cell lung cancer extracellular vesicles. Cell Biosci. 2018; 8: 2.

40. Wang T, Jian Z, Baskys A, Yang J, Li J, Guo H, Hei Y, Xian P, He Z, Li Z, Li N, Long Q. MSC-derived exosomes protect against oxidative stress-induced skin injury via adaptive regulation of the NRF2 defense system. Biomaterials. 2020; 257: 120264.

41. Ren S, Chen J, Duscher D, Liu Y, Guo G, Kang Y, Xiong H, Zhan P, Wang Y, Wang C, Machens HG, Chen Z. Microvesicles from human adipose stem cells promote wound healing by optimizing cellular functions via AKT and ERK signaling pathways. Stem Cell Res Ther. 2019; 10: 47

