## Supplemental figure 1 and 2 for "Chronological aging impacts abundance, function and microRNA content of extracellular vesicles produced by human epidermal keratinocytes": Supplemental data.docx

Heatmap representing the expression of the 100 miRNAs with the highest variance in the 12 EV samples analyzed in our study. The miRNAs that are significantly (Padj<0.05) induced in old keratinocytes EVs compared to young keratinocytes EVs at early passage (P2) are indicated by a red box. The miRNAs that are significantly (Padj<0.05) repressed in old keratinocytes EVs compared to young keratinocytes EVs at early passage (P2) are indicated by a blue box.

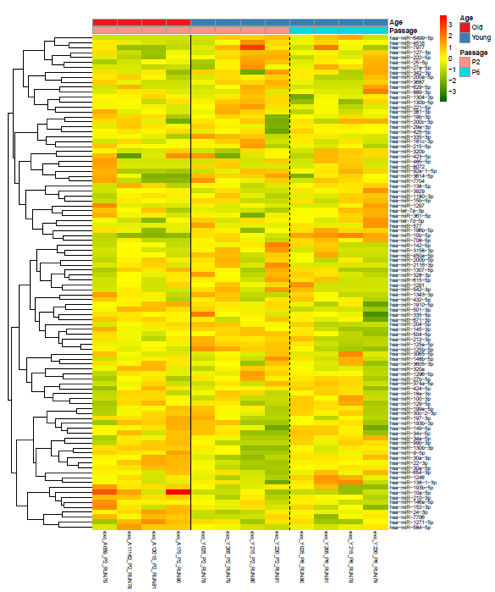

**Supplemental Figure 2**

Immunofluorescence labelling of Reconstructed Human Epidermis (RHE) obtained with old keratinocytes (A to H) and treated with young keratinocytes EVs (A to D) or obtained with young keratinocytes (I to P) and treated with old keratinocytes EVs (I to L). Keratin 1 (K1) (A, E, I, M) and involucrin (B, F, J, N) protein expressions were investigated. The dot line corresponds to the limit between the epidermal tissue and the substrate membrane.

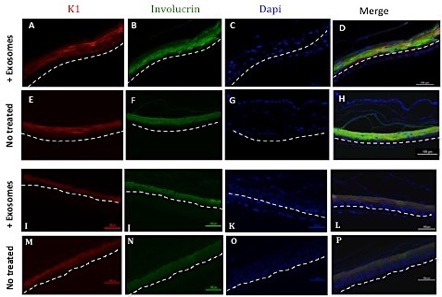
